# Genotype-phenotype correlations and de novo induction of cancer stem cells in Wilms tumor initiation

**DOI:** 10.1101/2025.11.19.689177

**Authors:** N.S. Pop, D. Koot, C.M. Brouwers, M.M. Linssen, J.W.C. Claassens, C.W.J. Cartlidge, D.D. Özdemir, K.S. Dolt, P. Hohenstein

**Affiliations:** Department of Human Genetics, Leiden University Medical Center, Leiden, Netherlands; Transgenic Facility Leiden, Central Animal Facility, Leiden University Medical Center, Leiden, Netherlands; Division of Developmental Biology, The Roslin Institute, Edinburgh, United Kingdom; European Research Institute for the Biology of Ageing, University of Groningen, University Medical Center Groningen, Groningen, The Netherlands; Koç University Research Center for Translational Medicine (KUTTAM), Istanbul, Turkey; Koç University School of Medicine, Istanbul, Turkey

## Abstract

Wilms tumor, the most common pediatric kidney cancer, arises from abnormal embryonic kidney development. Therapy resistance and tumor recurrence remain major challenges, likely driven by the presence of Cancer Stem Cells (CSCs). Here, we elucidate the earliest pathogenic events in genetically engineered mouse models exhibiting loss of *Wt1* or *LIN28B* overexpression, two Wilms tumor driver genes. Loss of *Wt1* leads to a disturbance of lineage identity of the mutant cells, whereas LIN28B leads to a disturbed transition between uninduced and induced NPC (nephron progenitor cell) state. In both cases the appearance of cells positive for all four Wilms tumor cancer stem cell markers is the result of the tumor initiating mutation. The existence of genotype-phenotype correlations in primary developmental phenotypes and cancer stem cell expression patterns has important implications for therapeutic opportunities and requirements.

## Introduction

Wilms tumor is the most common pediatric renal malignancy, occurring in approximately 1 in 10,000 children, typically before the age of five (Spreafico et al., 2021). They arise from aberrant embryonic kidney development, potentially during the critical window of definitive kidney development between the fifth and eight week of gestation (Zheng et al., 2023). Standardized diagnostic and therapeutic procedures have made it possible to cure nearly 90% of the affected children. However, therapy resistance, postoperative recurrence, and high-risk Wilms tumors remain significant clinical challenges. Histological analyses consistently reveal persistence of embryonic renal structures within tumors (Beckwith et al., 1990; Hohenstein et al., 2015), including a subset of tumor cells resembling progenitor-like cells. These observations are consistent with the cancer stem cell (CSC) model, in which a small population of stem cell-like cells drives tumor initiation, recurrence, and therapy resistance, potentially explaining the poor outcomes observed in high-risk Wilms tumor (Bonnet and Dick, 1997; Ginestier et al., 2007; Pode-Shakked et al., 2013). Understanding the origin of these Wilms tumor CSCs would provide valuable insights in the biology of WT and the earliest pathogenic events driving Wilms tumor formation, thus providing new leads to improve patient prognosis and develop targeted therapies.

Several studies defined the Wilms tumor CSC in patient-derived material (Petrosyan et al., 2023; Pode-Shakked et al., 2013). NCAM1^+^ cells with high ALDH activity were found capable of initiating xenograft tumors, fulfilling key CSC criteria, although these cells lost tumorigenic potential when cultured *ex vivo* (Pode-Shakked et al., 2009; Pode-Shakked et al., 2013). More recently a SIX2^+^CITED1^+^ population was described as a different kind of CSC, maintaining tumor-initiating capacity in culture and meeting the CSC criteria (Petrosyan et al., 2023).

These identified CSC markers have distinct functions during normal embryonic kidney development. Polysialylated NCAM1 (PSA-NCAM1), the embryonic version of the neural cell adhesion molecule 1 (NCAM1), is an important protein that mediates cell-cell adhesion, is expressed on NPCs and endothelial cells and facilitates cell movement during kidney development (Niculovic et al., 2025). The protein is expressed in undifferentiated metanephric mesenchyme at E11.5-E12.5 and in early nephron structures (comma- and S-shaped bodies) during kidney development, but is absent postnatally (Markovic-Lipkovski et al., 2007). *Ncam1*-deficient mice exhibit impaired glomerular vascularization, with reduced endothelial cell numbers and smaller glomeruli, indicating an essential role for NCAM1 in kidney vascularization in the mouse embryonic kidney (Niculovic et al., 2025). ALDH activity, measurable using the ALDEFLUOR assay, is a widely used CSC marker (Ginestier et al., 2007). Multiple ALDH isoforms contribute to this signal, including ALDH1A1, ALDH1A2, ALDH1A3 and ALDH2 (Zhou et al., 2019). ALDH enzymes synthesize retinoic acid from retinaldehyde, a critical signaling molecule that regulates cell differentiation, proliferation, and tissue patterning (Niederreither et al., 1999). Expression is lineage- and stage-specific: *Aldh1a1* and *Aldh1a3* are dynamically expressed in differentiating tubular structures derived from the nephrogenic lineage or ureteric bud, whereas *Aldh1a2* is mainly localized in the cortical stroma and essential for ureteric bud outgrowth and nephron formation, and is weakly expressed in the early nephron (comma- and S-shaped bodies, and in the glomerulus) (Li et al., 2020; Li et al., 2017).

Transcription factors *Six2* and *Cited1* mark uninduced NCPs (Kobayashi et al., 2008; Mugford et al., 2009). Upon NPCs induction, *Cited1* expression is lost first, followed by *Six2* (Mugford et al., 2009). The persistence of SIX2^+^CITED1^+^ populations in Wilms tumor xenografts suggests that these tumors may originate from nephron progenitors that failed to undergo proper differentiation, although this developmental process has yet to be conclusively demonstrated. The normal expression pattern of these CSC markers suggest that NCAM1^+^ALDH-activity is a natural occurrence in early nephron structures (comma- and S-shaped bodies), whereas SIX2^+^CITED1^+^ reflects a transcriptional signature of uninduced NPCs.

To date, 46 Wilms tumor predisposition and driver genes have been identified, many of which are essential for embryonic kidney development and are classified in different functional groups (Spreafico et al., 2021). Most existing data on biological aspects of Wilms tumors was obtained without knowing the mutational status of Wilms tumor genes in the samples and it is therefore unclear how universally applicable the findings are for any given Wilms tumor. The different functional classifications of Wilms tumor genes might lead to different underlying biology of tumors with different mutations, and if so, different opportunities or requirements for therapeutic intervention. It will also affect the outcome of clinical trials if the patients are not properly matched to the treatment that is being tested. Even if full genome sequencing of Wilms tumor samples would become routine, without understanding what biological effects different mutations have this will remain a major obstacle to further improving Wilms tumor patient outcome.

Wilms Tumor 1 (*WT1*) is the most extensively studied Wilms tumor gene. Although best known as a transcription factor, WT1 can bind both RNA and DNA, functioning as either an activator or repressor depending on the cellular and chromatin context or in RNA metabolism (Essafi et al., 2011; Hohenstein and Hastie, 2006). Although best known as a transcription factor, WT1 can bind both RNA and DNA, functioning as either an activator or repressor depending on the cellular and chromatin context or in RNA metabolism (Essafi et al., 2011; Hohenstein and Hastie, 2006). During nephrogenesis, WT1 is critical for driving mesenchymal-to-epithelial transition (MET) in NPCs. Approximately 15% of patients carry loss-of-function *WT1* mutations, which are frequently associated with ectopic tissue differentiation into skeletal muscle, fat, and less commonly bone or cartilage (Miyagawa et al., 1998; Schumacher et al., 2003).

Another Wilms tumor driver gene, *LIN28B*, is primarily expressed in progenitor cells, where it promotes proliferation and blocks differentiation by inhibiting maturation of the let-7 microRNAs, thereby upregulating oncogenic targets including *RAS*, *MYC*, and *HMGA2* (Viswanathan and Daley, 2010). LIN28B has been implicated in tumor progression, epithelial-to-mesenchymal transition, CSC formation, and therapy resistance (Cotino-Najera et al., 2024; Li et al., 2019; Zhou et al., 2013). Its precise role in kidney development is unclear, its declining expression by embryonic day (E)16.5 suggests a temporally restricted function during nephrogenesis (Viswanathan and Daley, 2010; Yermalovich et al., 2019). The earliest phenotypic events following *WT1* loss or *LIN28B* overexpression in Wilms tumor patients remain largely unknown, limiting our understanding of how these mutations initiate tumorigenesis.

Genetically engineered mouse models provide a powerful model to study the earliest phenotypic effects of tumor-initiating mutations in a controlled developmental context. These models allow us to study the direct consequences of Wilms tumor-initiating mutations and track emergence, expansion, and expression of cell populations that could act as CSCs. Here we use lineage specific loss of *Wt1* and *LIN28B* overexpression mouse models to study the genotype-phenotype correlations during kidney development, and their effects on CSC marker expression.

## Results

### Conditional Wt1 knockout models

We previously knocked out *Wt1* at different stages of nephron development using *Nes*-Cre, *Pax8*-Cre a *Wnt4*-Cre drivers, showing that the earliest, pre-MET loss of *Wt1* using *Nes*-Cre most closely resembled *WT1*-mutant tumors. It led to an expansion of mesenchyme and a block of MET (Berry et al., 2015). The metanephric mesenchyme gives rise to the condensed mesenchyme containing the nephron progenitor cells (NPCs), expressing *Six2* (Kobayashi et al., 2008), and the remaining uncondensed mesenchyme containing stromal progenitor cells with a distinct expression pattern of *Foxd1* (Kobayashi et al., 2014). Both populations transiently express *Wt1* (Armstrong et al., 1993; Berry et al., 2015; Weiss et al., 2020). The *Nes*-Cre model affects both lineages, and therefore cannot distinguish between the roles of *Wt1* in them. To determine the nephrogenic- and stromal-specific effects when *Wt1* is lost, we used three Cre-drivers: *Six2*^GFPCre-IRES-PuroR^ (*Six2*^GCiP^) to target the nephrogenic lineage, *Foxd1*^GFPCre^ (*Foxd1*^GC^) to target the stromal lineage, and *Wt1*^CreERT2^ (*Wt1*^CE^) for targeting both lineages in a tamoxifen-dependent controlled manner (Dolt et al., 2013; Kobayashi et al., 2014; Zhou et al., 2008). We crossed the three Cre-drivers with the *Wt1* conditional allele (Martinez-Estrada et al., 2010) and in some cases included the lox-stop-lox tdTomato reporter (Madisen et al., 2010) to follow the lineage of mutant and control cells. This conditional *Wt1* knockout model is a phenocopy of the conventional model (Kreidberg et al., 1993) when driven by a germline Cre deleter strain, and does not have a phenotype when driven by the *Hoxb7*-Cre ureteric bud-specific driver (Berry et al., 2015).

### *Six2*-driven Wt1 loss blocks nephrogenesis

To selectively delete *Wt1* in the nephrogenic lineage, we used the *Six2*^GCiP^ driver to knockout *Wt1* in the developing kidney (Fig. 1A). *Six2* is expressed in NPCs from E10.5 in the metanephric mesenchyme and later in the cap mesenchyme, and is predicted to be a direct transcriptional target of *Wt1* (Kobayashi et al., 2008; O’Brien et al., 2018). As NPCs form pretubular aggregates (PTA), which epithelializes into a renal vesicles (RV), comma-shaped and S-shaped bodies to form a mature nephron (McMahon, 2016), *Six2* expression is lost.

**Figure 1:**
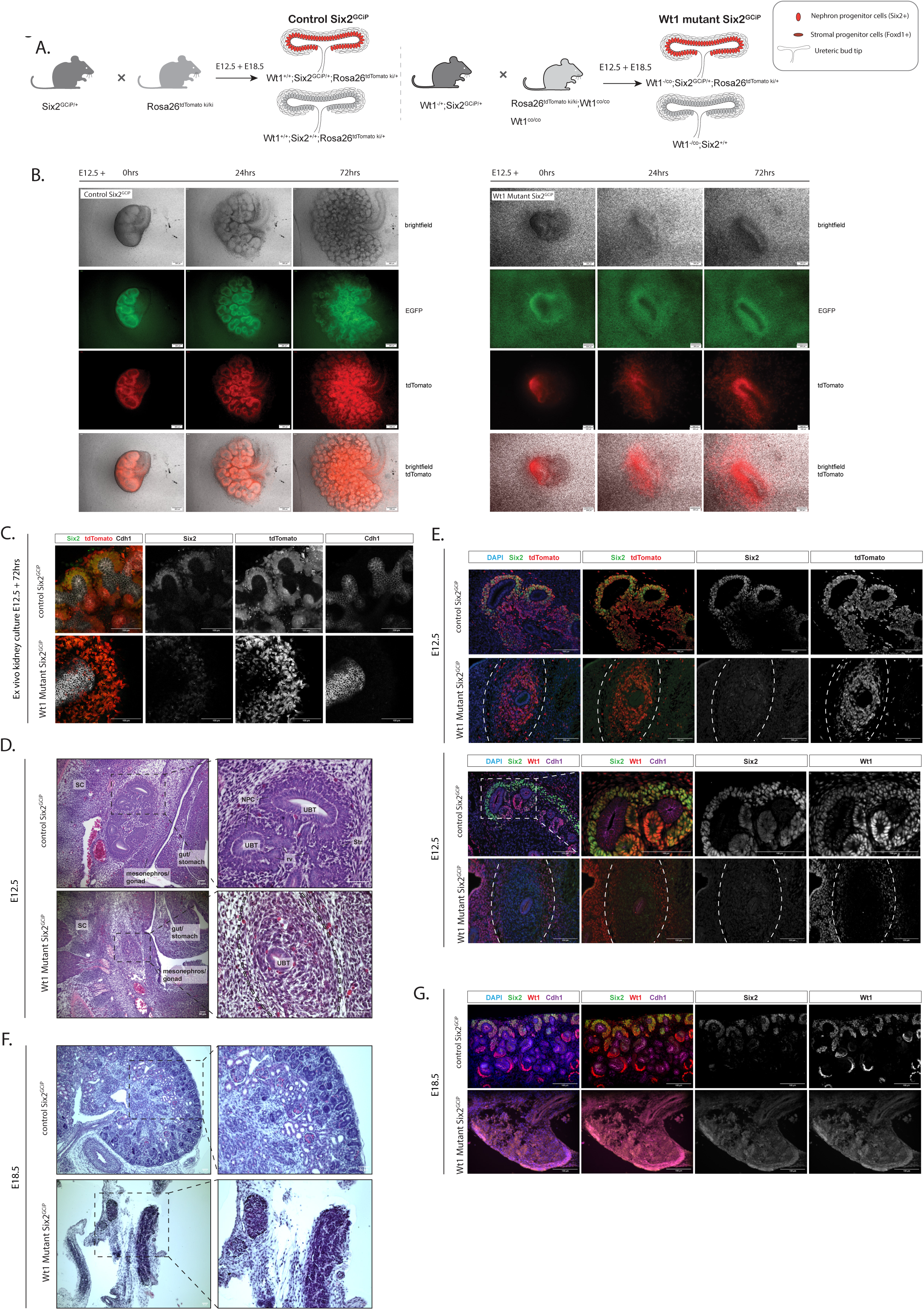
*Six2*^GCiP^ driven *Wt1*-mutant model characterization. (A) Schematic representation of the *Six2*^GCiP^ lineage specific knockout of *Wt1* in the mouse model. (B) *Ex vivo* kidney organ culture of E12.5 control (left) and *Wt1* mutant kidney (right), with eGFP linked to SIX2 expression and TdTomato representing the lineage. Scale 200µm. (C) Staining for SIX2, TdTomato and CDH1 on *ex vivo* kidney organ cultures of a control and *Wt1* mutant kidney. Scale 100µm. (D) H&E staining of control and *Wt1* mutant kidneys at E12.5. Scale 50µm. (E) Staining for TdTomato to validate lineage and WT1 to validate knockout in E12.5 control and mutant kidneys. Scale 100µm. (F) H&E staining of control and *Wt1* mutant kidneys at E18.5. Scale 50µm. (G) Staining for SIX2, CDH1 and WT1 to elucidate identity of E18.5 mutant kidneys. Scale 100µm.

*Ex vivo* kidney organ culture time lapse imaging of the *Six2*^GCiP^ allele (Dolt et al., 2013) in control E12.5 embryonic kidneys confirmed *Six2*-linked GFP expression in the cap mesenchyme, and the tdTomato lineage reporter confirmed contribution to the nephrogenic lineage, with exclusion from the stroma and ureteric bud (Fig. 1B, left panel; supplementary movie 1A). In contrast, e*x vivo* kidney organ cultures of E12.5 *Six2*^GCiP^ driven *Wt1*-mutant kidneys showed severely disrupted nephron differentiation and ureteric bud branching morphogenesis (Fig. 1B, right panel; supplementary movie 1B). At 0hrs, the *Wt1*-mutant eGFP expressing cells surrounded the ureteric bud tip, but subsequently the eGFP signal decreased, indicative of cells losing SIX2-expression, consistent with *Six2* as potential *Wt1* target (O’Brien et al., 2018). The tdTomato-expressing mutant cells accumulated around the ureteric bud tip, or migrated rapidly outwards. Whole mount immunohistochemical staining confirmed loss of SIX2 in the tdTomato-expressing cells (Fig. 1C).

Consistent with the time lapse imaging, histological analysis of E12.5 embryos showed ureteric bud tips surrounded by mesenchymal-like cells (Fig. 1D). Immunohistochemistry confirmed loss of SIX2 expression in tdTomato-expressing cells (Fig. 1E, top panels), which had lost WT1 expression and surrounded a CDH1-positive tubular structure (Fig. 1E, bottom panels). Surrounding stroma, negative for tdTomato, continued to express WT1. Proliferative marker KI67 is reduced in the kidney region (Sup. Fig. 5A-B).

At E18.5, mutants exhibited structures resembling early budding stage. Sectioning the isolated urogenital system complicated the identification of the kidney remnants in 2D, due to their lower localization in the embryo (Fig. 1F, arrows). The derived structures were negative for SIX2, WT1, and CDH1 expression, making it difficult to define their developmental stage, cellular identity, or structural nature. Nevertheless, these structures were located in the anatomical region where kidneys are expected to form.

### *Foxd1*-driven Wt1 loss disrupts ureteric bud branching and cortical organization

We used the *Foxd1*^GC^ driver (Kobayashi et al., 2014) to delete *Wt1* specifically in FOXD1^+^ stromal progenitors (Fig. 2A). The FOXD1^+^ stromal progenitors originate from the uncondensed mesenchyme and are first present around E11.0, when they begin populating the cortical and medullary interstitium, the fibrocytes of the capsular tissue and the vascular-supporting pericytes (Kobayashi et al., 2014).

**Figure 2:**
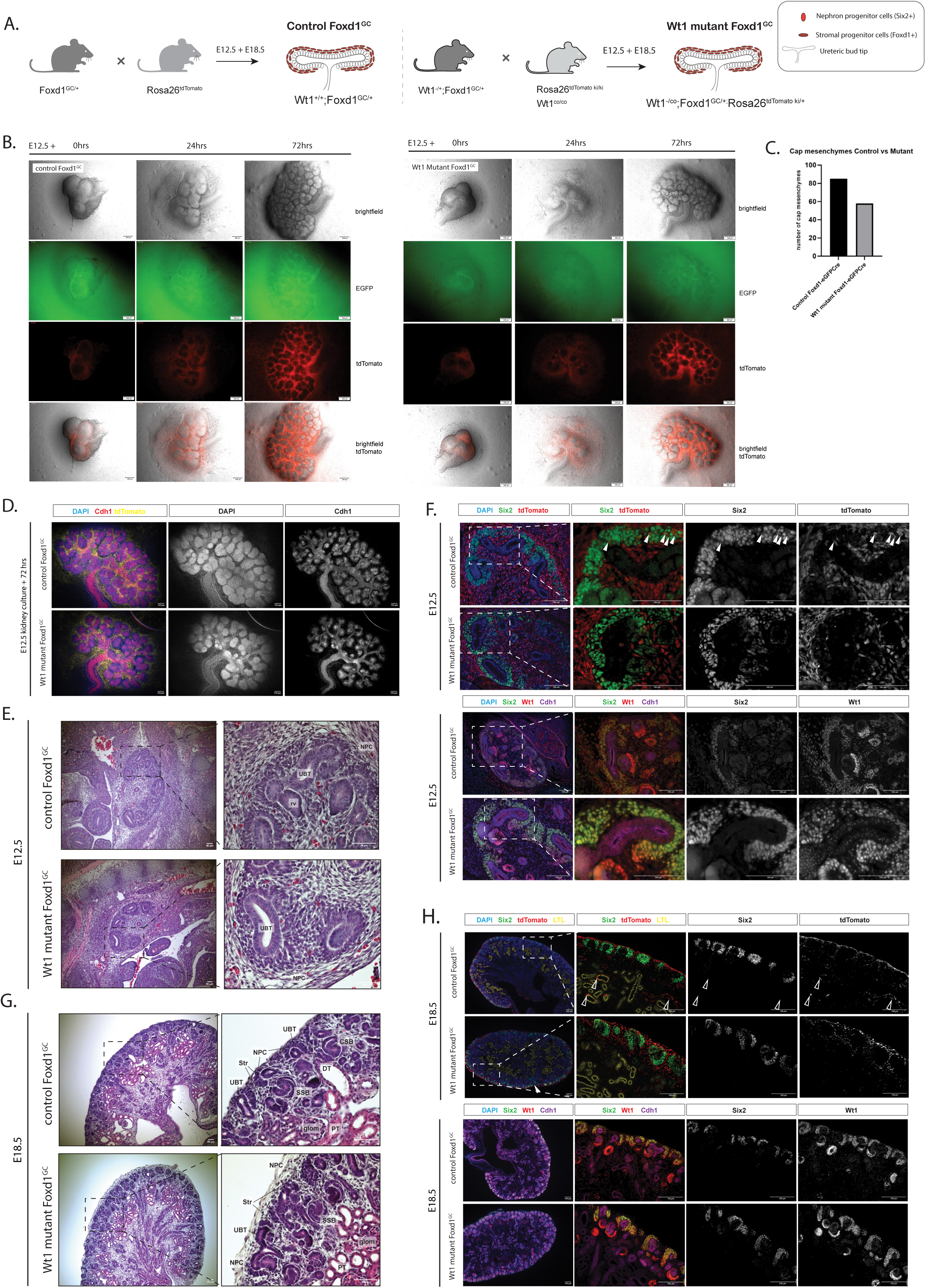
*Foxd1^GC^* driven *Wt1*-mutant model characterization. (A) Schematic representation of the *Foxd1*^GC^ lineage specific knockout of *Wt1* in the mouse model. (B) *Ex vivo* kidney organ culture of E12.5 control (left) and *Wt1* mutant kidney (right), with eGFP linked to FOXD1 expression and TdTomato representing the lineage. Scale 200µm. (C) Absolute quantification of the number of cap mesenchymes in the control versus mutant kidney, quantified from the *ex vivo* kidney organ culture. (D) Staining of DAPI and CDH1 on the *ex vivo* kidney culture. Scale 100µm. (E) H&E staining of control and *Wt1* mutant kidneys at E12.5. Scale 50µm. (F) Staining for TdTomato to validate lineage and WT1 to validate knockout in E12.5 control and mutant kidneys. Scale 100µm. White arrowheads indicate Foxd1 lineage contribution to SIX2+ cells. (G) H&E staining of control and *Wt1* mutant kidneys at E18.5. Scale 50µm. (H) Staining for TdTomato to validate lineage and WT1 to validate knockout in E18.5 control and mutant kidneys. Scale 100µm. White open arrows indicate Foxd1-lineage in or around proximal tubules.

Lineage tracing of *Foxd1*^GC^ through tdTomato expression in *ex vivo* control kidney organ cultures confirmed expression in the stromal lineage surrounding the nephrogenic caps and branched ureteric bud (Fig. 2B, left panel; supplementary movie 2A). In contrast, *ex vivo* organ cultures of *Foxd1^GC^* driven *Wt1*-mutant kidneys showed altered ureteric bud morphogenesis, characterized by reduced branching complexity, elongated and sparser branches, increased spacing between them, and fewer terminal tips (Fig. 2B-C; supplementary movie 2B) - a phenotype similar to that described previously (Weiss et al., 2020). Mutant kidneys are smaller, consistent with reduced branching complexity and fewer nephrogenic caps (Fig. 2D).

Histological analysis of E12.5 sections confirmed decreased ureteric bud branching in the *Wt1*-mutants (Fig. 2E). Ureteric bud tips were surrounded by loosely organized NPCs. The *Foxd1*-lineage confirmed strong contribution to the stromal compartment (Fig. 2F, upper panels). We observed some SIX2^+^ tdTomato^+^ cells in the cap mesenchyme of control kidneys (Fig. 2F, white arrows), though these cells expressed lower SIX2 levels compared to surrounding cap mesenchymal. Such double-positive cells were absent in *Foxd1*-driven *Wt1*-mutant kidneys. Antibody staining confirmed loss of WT1 in the stroma surrounding the cap mesenchyme, while the cap mesenchyme itself remained WT1^+^ (Fig 2F lower panels).

At E18.5, mutant kidneys appeared smaller, with expanded cortical stroma between the developing nephrons – consistent with the phenotype described by Weiss et al (Fig. 2G). We observed disrupted nephrogenic zone organization, including SIX2^+^ caps that lost their apical orientation and started moving inwards (Fig. 2H upper panels, white filled arrow; Sup. Fig. S1). This may be the consequence of expansion and migration of cortical stromal cells into the nephrogenic zone. We detected tdTomato^+^ cells in or around proximal tubules (LTL^+^) (Fig. 2H, white open arrows). It is not clear if these cells are part of the proximal tubules. Staining for WT1 in control kidneys at E18.5 showed low expression in the stroma, which was absent in mutants (Fig. 2H lower panels). Proliferative marker KI67 is highly expressed in the nephrogenic zone, comparable to the control kidneys (Sup. Fig. 5A-B).

### Combined nephrogenic and stromal Wt1 loss delays kidney development and initiates blastemal features

Targeting the nephrogenic lineage alone caused a severe developmental block, whereas targeting the stromal lineage led to stromal overgrowth, nephrogenic delay, and altered ureteric bud morphogenesis, but each was different form the *Nes*-Cre driven *Wt1* knockout (Berry et al., 2015). Since *Wt1* is expressed in both lineages we combined the conditional *Wt1* knockout and tdTomato reporter alleles with the *Wt1*^CE^ knock-in driver (Zhou et al., 2008). The latter is itself a *Wt1*-inactivating allele, so in the tamoxifen induced embryos the Cre-induced loss of the conditional *Wt1* copy results in complete loss of the protein (Fig. 3A). To determine the best window and dosage for targeting the SIX2^+^ nephrogenic lineage and surrounding stroma, we tested single dose Tamoxifen (Tx) injections at E8.5, E9.5, E10.5 or E11.5, on embryonic reporter kidneys isolated at E12.5 and *ex vivo* cultured the kidneys from the treated litters for 48-72hrs. The tdTomato signal indicated the lineage of cells where recombination happened (Fig. 3B). Single Tx injections targeted most of the cells of both lineages when injected at E9.5 or E10.5. Double injections at E9.5 and E10.5 increased this number of targeted cells in the nephrogenic zone. Immunostaining for SIX2 expression showed some cells that lacked tdTomato expression, indicating incomplete Cre activity (Fig. 3B, white arrows). This observation of “escaping”-cells is similar to what has been described for the *Nes*-Cre driven *Wt1* conditional knockout model (Berry et al., 2015) although in the current case it is likely due to incomplete activation of the recombinase by tamoxifen.

**Figure 3:**
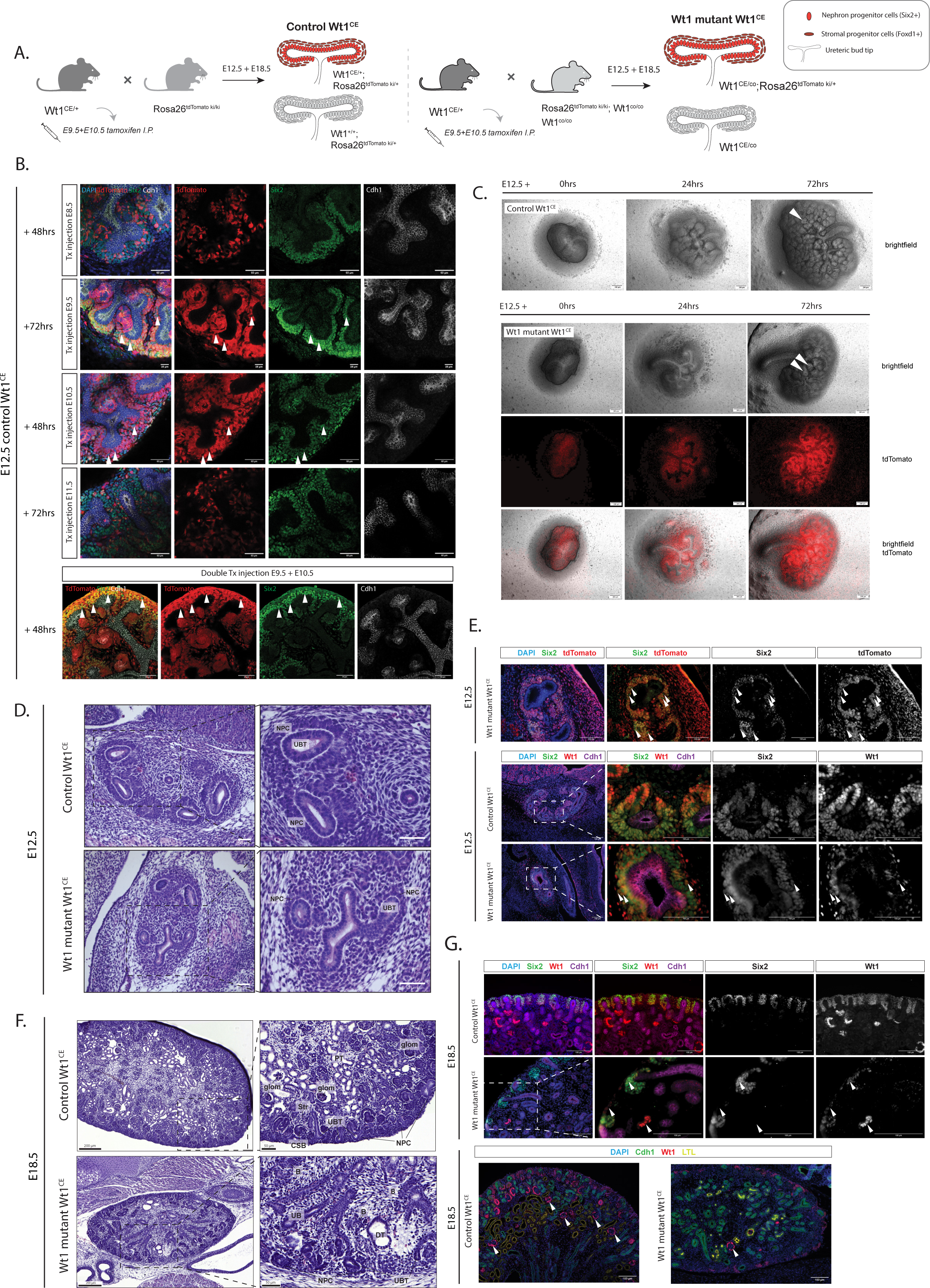
*Wt1^CE^* driven *Wt1*-mutant model characterization. (A) Schematic representation of the *Wt1^CE^* spatial and temporal specific knockout of *Wt1* in the mouse model. (B) Optimization of Tx injection at E8.5, E9.5, E10.5 or E11.5, and double injections at E9.5 + E10.5, analyzed in *ex vivo* kidney cultures at E12. Cells labeled with white arrows are a representation of the tamoxifen treatment escaping cells. White arrowheads indicate cap mesenchyme cells that escaped tamoxifen-induced Cre activation. Scale 60µm. (C) *Ex vivo* kidney organ culture of E12.5 control (upper) and *Wt1* mutant kidney (lower), with TdTomato representing the lineage. White arrows highlight epithelializing structures. White arrowheads indicate epithelialized structures based on morphology of the cells. Scale 200µm. (D) H&E staining of control and *Wt1* mutant kidneys at E12.5. Scale 50µm. (E) Staining for TdTomato to validate lineage and WT1 to validate knockout in E12.5 control and mutant kidneys. White arrowheads indicate cap mesenchyme cells that escaped tamoxifen-induced Cre activation. Scale 100µm. (F) H&E staining of control and *Wt1* mutant kidneys at E18.5. Scale 50µm. (G) Staining for TdTomato to validate lineage and WT1 to validate knockout in E18.5 control and mutant kidneys. LTL marks proximal tubules. White arrows indicate nephrogenic(-derived) WT1 expression indicating the escaped regions. Scale 100µm.

We used the double E9.5+E10.5 Tx treatment to knockout *Wt1*. We compared mutant kidneys to littermate controls which lacked the CreERT2 allele, but were equally exposed to Tx. In this case the control samples lack Cre activity and will therefore not activate the tdTomato Cre reporter. Lineage tracing of *Wt1*-mutant kidneys revealed smaller kidneys with altered ureteric bud branching and fewer nephrogenic caps (Fig. 3C; supplementary movie 3A-B), similar to the *Foxd1*^GC^ driven mutants. The morphology of the cells suggested that mutant cap mesenchymes lacked epithelialization (Fig. 3C, white arrows), a sign of delayed or blocked MET, a process in which we showed WT1 plays a key role before (Berry et al., 2015; Davies et al., 2004; Essafi et al., 2011).

Histological assessment at E12.5 showed a thinner NPC layer in mutants compared to controls (Fig. 3D). The NPC population is dispersed, lacking the tight cap-like structure around the ureteric bud tips. The ureteric bud epithelium appears less branched compared to the control. The stroma in the control is relatively thin and confined around the nephrogenic zone, where in the mutant it appears to be expanded – more stroma surrounding the ureteric bud tips and nephron progenitors. The stroma is less organized and seems loosely packed.

Immunohistochemistry for SIX2 and tdTomato confirmed incomplete Cre activation in mutants, evidenced by SIX2^+^ tdTomato^-^ cells (Fig. 3E, white arrows), while controls lacked CreERT2 and were tdTomato^-^ (data not shown). Co-staining for SIX2 and WT1 revealed double-positive cells, the “escaping” SIX2^+^ tdTomato^-^ cells (Fig. 3E lower panels). CDH1 staining further supported the presence of widened, misshaped epithelium (Fig. 3E lower panels).

Mutant kidneys at E18.5 were markedly smaller, lacked clear corticomedullary segmentation, and displayed severely altered nephrogenesis, with only a few glomeruli present (Fig. 3F). Embryonic-like epithelium was observed in the nephrogenic zone, with emerging blastemal-like regions (Fig. 3F). Staining for proliferative mark KI67 reveals disorganized expression of the protein, with high expression observed in the cortical-region (Sup. Fig. 5B white arrows). Stromal overgrowth was also evident. WT1 immunohistochemistry revealed “escaping” nephrons (Fig. 3G), consistent with observations at E12.5. Mutant kidneys displayed reduced LTL^+^ proximal tubules, consistent with impairment of MET and blocked or delayed nephron differentiation (Fig. 3G).

ALDH1A2 is the primary contributor to ALDEFLUOR activity in Wilms tumour CSCs

Wilms tumor blastema is associated with the Wilms tumor Cancer Stem Cells (CSCs). As mentioned above, the two sets of Wilms tumor CSC markers (NCAM1^+^ with ALDEFLUOR-activity (Pode-Shakked et al., 2013), and SIX2^+^CITED1^+^ (Murphy et al., 2012; Petrosyan et al., 2023)) are both linked to tumor maintenance and progression. In the first marker set, ALDH activity was measured using the ALDEFLUOR assay, which requires living cells, analyzed through flow cytometry. To investigate whether similar NCAM1/ALDH patterns occur during kidney development, we repeated the NCAM1/ALDERED assay (analogous to ALDEFLUOR) on embryonic mouse kidneys. No significant differences were observed between mutant and control kidneys for all lineage-specific *Wt1* mutant kidneys (Sup. Fig. 2).

To visualize the protein expression levels of the CSC markers over the course of embryonic kidney development, we first determined which ALDH-paralogue was the primary contributor to the ALDEFLUOR assay in WT patient samples. ALDH1A1, ALDH1A2, ALDH1A3 and ALDH2 can all contribute to the detected signal (Zhou et al., 2019). In the context of Wilms tumor, ALDH1A2 has been suggested to be the paralogue associated with CSCs (Petrosyan et al., 2023; Wegert et al., 2020), but its specificity for Wilms tumors has not yet been proven. We reanalyzed microarray data (Shukrun et al., 2014) from Wilms tumor xenograft fractions unsorted and sorted for NCAM1^+^ALDEFLUOR^+^ and NCAM1^+^ALDEFLUOR^-^ (Sup. Fig. 3A). The NCAM1+/ALDEFLUOR+ fraction shows high expression for both *ALDH1A2* and *ALDH1A3*. Nonetheless, *ALDH1A3* expression is not enriched in the CSC fraction given its higher expression in the unsorted fraction. *ALDH1A1* is low in all fractions and *ALDH2* is high in unsorted and decreased in both sorted fractions. *ALDH1A2* is the only paralogue specifically upregulated in the NCAM1+/ALDEFLUOR+ faction and therefore these results indicate that ALDH1A2 is the principal contributor to the ALDEFLUOR activity in these Wilms tumor CSCs.

To examine ALDH1A-paralogue expression in the developing mouse kidney, E12.5 embryonic kidneys were cultured *ex vivo* for 24, 48 and 72 hours and stained for ALDH1A1, ALDH1A2, and ALDH1A3. ALDH1A1and ALDH1A3 were weakly detected in the ureteric bud epithelium, whereas ALDH1A2 showed strong expression in the cortical region of the kidney, in close proximity of the nephrogenic caps, presumably marking stromal cells (Sup. Fig. 3B). Only in the E12.5 kidneys cultured for 48hrs weak ALDH1A2 expression was observed in renal vesicle-shaped structures. Co-staining with FOXD1 antibodies confirmed the stromal identity of the ALDH1A2 positive cells. These findings support ALDH1A2 is mainly localized to the cortical stroma, with weaker protein expression patterns in the epithelializing nephron (Sup. Fig. 3C).

### CSC Marker sets for committed and uncommitted NPCs

The CSC marker sets were identified on xenograft derived tumor tissues (Petrosyan et al., 2023; Pode-Shakked et al., 2013), which might not reflect the developmental stage of origin anymore. To map these CSC markers onto normal developing kidneys and examine their dynamics in the context of Wilms tumor mutations, we performed immunohistochemistry for NCAM1, ALDH1A2, SIX2, and CITED1 in embryonic kidneys from control and *Wt1*-mutants at E12.5 and E18.5. Due to overlapping antibody host species, ALDH1A2 and CITED1 were analyzed on consecutive sections, resulting in two staining combinations: (NCAM1, ALDH1A2, SIX2; and NCAM1, CITED1, SIX2.

In normal embryonic kidneys, NCAM1^+^ALDH1A2^+^SIX2^+^ triple-positive cells localized to the SIX2^+^CITED1^-^ committed NPC/PTA population (Fig. 4A, upper panels, white arrows). These committed NPC/PTA cells exhibited low ALDH1A2 expression compared to surrounding stromal cells. In contrast, the uninduced SIX2^+^CITED1^+^ cells co-express NCAM1, while they lack ALDH1A2 expression, defining a more uninduced NCAM1^+^CITED1^+^SIX2^+^ population (Fig. 4 A, lower panels). Therefore, in normal developing kidneys ALDH1A2 and CITED1 display a mutually exclusive expression pattern. Early nephron structures such as comma- and S-shaped bodies strongly expressed NCAM1 and ALDH1A2 but lacked SIX2 and CITED1, consistent with expression patterns in other studies (Li et al., 2020; Li et al., 2017; Markovic-Lipkovski et al., 2007).

**Figure 4:**
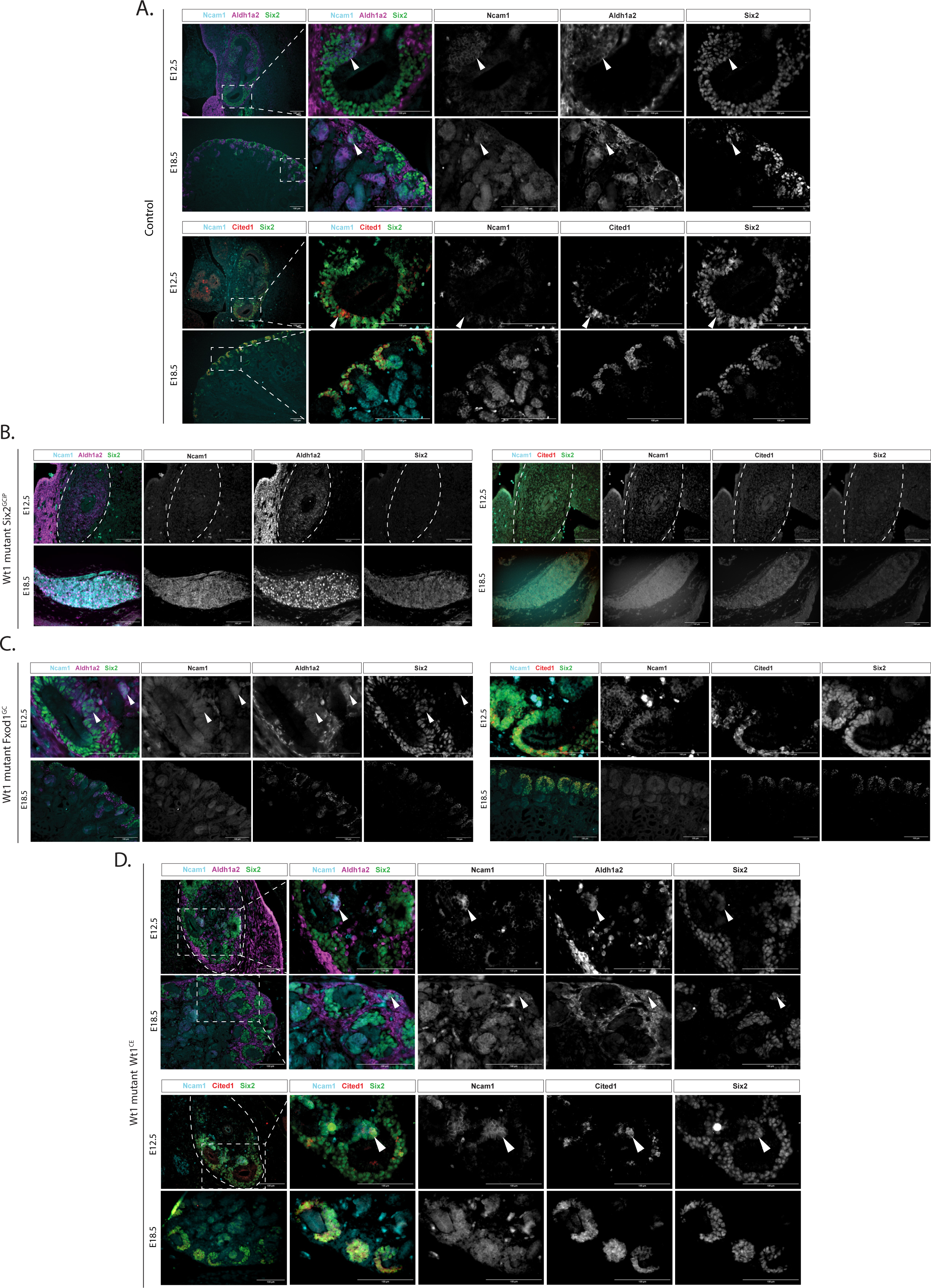
Cancer Stem Cell marker expression in lineage-specific *Wt1* mutant kidneys. (A) Staining for CSC markers NCAM1, ALDH1A2, SIX2 and CITED1 in the normal developing embryonic kidney at E12.5 and E18.5. White arrows indicate triple-marker expressing cells. Scale 100µm. (B) CSC marker expression in *Six2^GCiP^* driven *Wt1* mutant kidneys at E12.5 and E18.5. Scale 100µm. (C) CSC marker expression in *Foxd1*^GC^ driven *Wt1* mutant kidneys at E12.5 and E18.5. White arrows indicate triple-marker expressing cells. Scale 100µm. (D) CSC marker expression in *Wt1*^CE^ driven *Wt1* mutant kidneys at E12.5 and E18.5. White arrows indicate triple-marker expressing cells. Scale 100µm.

Lineage-specific *Wt1* loss revealed distinct effects on these populations. *Six2*^GCiP^-driven *Wt1* mutants at E12.5 showed a reduction in NCAM1, CITED1, and SIX2 expression, while ALDH1A2 expression was observed in the same region where *Wt1* was lost (Fig. 4B upper panels, Fig. 2E). This suggests that *Wt1*-deficient cells obtained an alternative identity, possibly differentiating towards stroma which is characterized by ALDH1A2 expression but not the other markers. At E18.5, the kidney remnants showed a distinct expression pattern as observed at E12.5. The kidney remnant represents a homogenous tissue structure where NCAM1 is expressed, ALDH1A2 is high in the nucleus, SIX2 is low and CITED1 is lowly expressed in the nucleus (Fig. 4B, lower panels). *Foxd1*^GC^ driven *Wt1* mutants resembled controls, without changed CSC marker expression at E12.5 (Fig. 4C). However, we do observe a decline in ALDH1A2 expression at E18.5 (Fig. 4C). In the *Wt1*^CE^ mutant model, at E12.5 NCAM1^+^ALDH1A2^+^SIX2^+^ triple-positive cells localized to the committed NPCs/PTA similar to controls (Fig. 4D upper panel). By E18.5, ALDH1A2^+^ regions expanded substantially, likely reflecting increased stromal content, whereas NCAM1^+^ALDH1A2^+^SIX2^+^ triple-positive committed NPCs/PTA were scarce, consistent with impaired MET (Fig. 4D). The NCAM1^+^CITED1^+^SIX2^+^ uninduced NPC population was present at E12.5, with CITED1 also observed in the committed NPCs/PTA (Fig. 4D lower panels, white arrow), demonstrating CITED1 retention beyond the uncommitted NPC stage. We cannot test directly if these triple-positive committed NPCs/PTA co-express ALDH1A2 due to the overlapping antibody species, but ALDH1A2 expression in similar structures was observed (Fig. 4D upper panels). By E18.5, the disorganized nephrogenic zone co-expressed NCAM1^+^CITED1^+^SIX2^+^ in the caps, in line with observations in the control kidneys (Fig. 4.D lower panel) representing ALDH1A2^-^ uninduced NPCs.

### LIN28B-overexpression induces developmental disorganization and early Wilms tumor blastema

To compare the lineage-specific *Wt1* loss phenotype and WT CSC marker behavior to other Wilms tumor mutations, we examined the effects of LIN28B overexpression on kidney development using the doxycycline (dox)-inducible model (Urbach et al., 2014). We used the same *Wt1^GC^* driver used by Urbach et al. to control lineage-specificity of the model, but added an additional *lgs7^TRE^*^-LSL-tdTomato^ allele (Madisen et al., 2015) to express a tdTomato reporter to identify lineage-specific dox-induced cells (Fig. 5A). Doxycycline was provided from beginning of gestation (E0). *Ex vivo* kidney organ cultures of control littermates lacked the *Wt1*^GC^ allele but were exposed to doxycycline throughout gestation and *ex vivo* culturing, and showed a normal developing embryonic kidney without the reporter because of the lack of the Cre allele (Fig. 5B, upper panel; supplementary movie 4A). Lineage tracing of the mutant kidneys in organ cultures confirmed dox induced tdTomato expression, thus LIN28B overexpression, in the three lineages of the developing kidney (Fig. 5B, lower panels). The tdTomato signal was particularly strong in the ureteric bud, while to lesser extend in the nephrogenic and stromal cells. Mutant kidneys in organ cultures exhibited enlarged nephrogenic caps and delayed epithelialization based on the morphology of the cells, identified at E12.5 + 72 hours of kidney culture (Fig. 5B; supplementary movie 4B).

Histological analysis of LIN28B mutants at E12.5 showed the first signs of delayed nephrogenesis with fewer renal vesicle-like structures and disorganized epithelium, and increased stroma between the caps (Fig. 5C, left panel). By E18.5, mutant kidneys displayed disorganized caps and surface indentations (Fig. 5C, right panel, orange arrows). The mutant kidney exhibited regions where normal nephrogenesis occurred, alternated with blastemal-like regions (Fig. 5C, red arrows and circles). These blastemal-like regions surrounding a tubular CDH1^+^ structure are possibly NPCs surrounding a ureteric bud tip that turned inwards, possibly explaining the observed indentations. Staining for SIX2 expression revealed that surface indentations are lined by SIX2^+^ caps turning inwards (Fig. 5D, lower panel). Morphologically this is suggestive for lobe formation, a phenomenon observed in human kidney development but not in mice. At E12.5, WT1 expression is low in the mutant compared to control. While at E18.5 the expression pattern is similar to control, but the caps are enlarged (Fig. 5D).

The lobes observed at E18.5 were lost by postnatal day 19 (P19). At P19, the cortico-medullary segmentation disappeared, and the abnormalities in mutant kidneys developed into regions where immature embryonal-appearing epithelium was accompanied by small areas of blastema and stroma, potentially representing beginning Wilms tumors (Fig. 5E). These blastemal-like regions co-expressed SIX2 and WT1, surrounding CDH1^+^ embryonal-appearing epithelium, possibly resembling embryonic cap mesenchymes (Fig. 5F). The SIX2^+^ blastemal-like regions exhibited expression patterns similar to committed NPCs, with gradual decline of SIX2 expression while WT1 remained robustly expressed (Fig. 5F, white arrows). The regions that lost SIX2 appeared to be capable of progressing into RV-like shapes (Fig. 5F, orange arrows).

**Figure 5:**
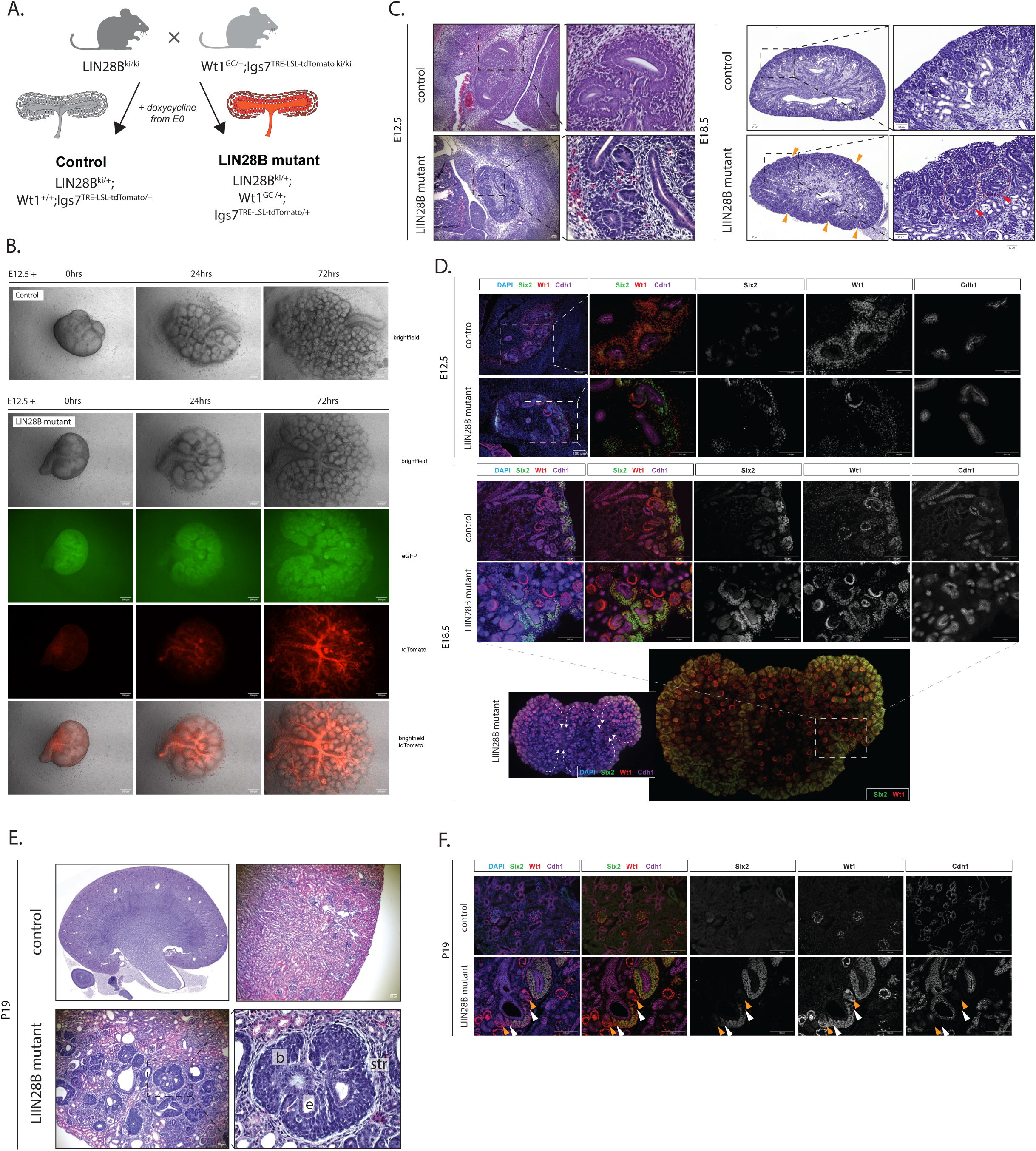
Characterization of the LIN28B overexpression model. (A) Schematic representation of the doxycycline inducible LIN28B overexpression model, with (representing mutants) and without (representing doxycycline exposed control littermates) tdTomato reporter. (B) *Ex vivo* kidney organ culture of E12.5 control (upper) and LIN28B mutant kidney (lower). Scale 200µm. (C) H&E staining of control and LIN28B mutant kidneys at E12.5 (left panel) and E18.5 (right panel). Scale 50µm. (D) Staining for WT1, SIX2 and CDH1 to identify phenotype at E12.5 and E18.5. Scale 100µm. (E) H&E staining of control and LIN28B mutant kidneys at P19. Scale 50µm. (F) Staining for WT1, SIX2 and CDH1 to identify blastemal phenotype at P19. White arrows indicate SIX2+WT1+ cells, orange arrows indicate SIX2-WT1+ cells, hinting towards commitment to the epithelial lineage. Scale 100µm.

### LIN28B-overexpression preserves embryonic marker profiles postnatally

We repeated the NCAM1/ALDERED assay on embryonic kidneys from the LIN28 model, but again no significant differences were observed between mutant and control kidneys (Sup. Fig. 4). Visualization of the four CSC protein markers revealed that the NCAM1^+^ALDH1A2^+^SIX2^+^ committed NPCs/PTA were absent in the mutant kidneys at E12.5, consistent with delayed nephrogenesis (Fig 6A upper panels). NCAM1^+^ALDH1A2^+^ epithelialized structures, similar to RV/comma- and S-shaped bodies, lacked SIX2 expression (Fig. 6A, white arrows). We observed NCAM1^+^CITED1^+^SIX2^+^ committed NPCs/RVs (Fig. 6A lower panels, white arrows), similar as described for the *Wt1*^CE^ *Wt1* mutant kidneys. By E18.5, NCAM1^+^ALDH1A2^+^SIX2^+^ triple-positive cells were detected within the nephrogenic zone in close proximity to the indentations (Fig. 6B, upper panel, white arrow). Their expression pattern resembles committed NPCs/PTA, but their spatial localization differed from that of canonical NPCs/PTAs within the nephrogenic niche. The observed positioning implies that these cells may have been captured in a state, potentially representing the earliest stages of blastema formation, rather than undergo nephron differentiation. This same blastemal-like population of cells co-expressed NCAM1^+^CITED1^+^SIX2^+^ in consecutive sections, suggesting all four CSC makers are co-expressed in the same cells in this early blastemal-like population (Fig. 6B lower panels, white arrows).

**Figure 6:**
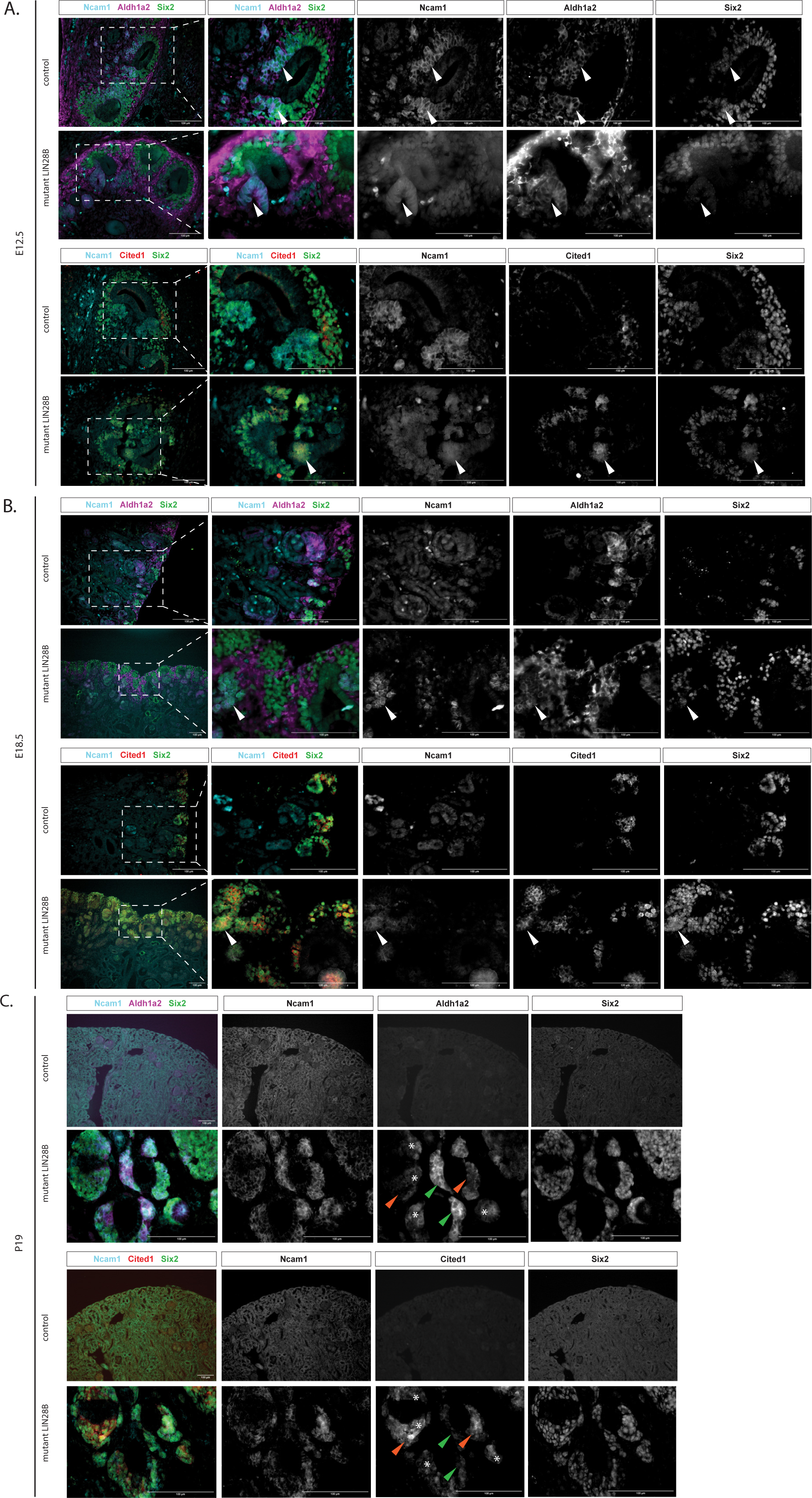
Cancer Stem Cell marker expression in LIN28B overexpression mutant kidneys. (A) CSC marker expression in E12.5 LIN28B overexpression kidneys. White arrows indicate triple-marker expressing cells. Scale 100µm. (B) CSC marker expression in E18.5 LIN28B overexpression kidneys, showing early blastemal-like cells co-expressing all four markers in the same cells indicated by the white arrows on consecutive sections. Scale 100µm. (C) CSC marker expression in P19 LIN28B overexpression kidneys, showing blastemal-like cells with distinct patterns for ALDH1A2 and CITED1 expression, and a transitional cell co-expressing all four markers in the same cell. High protein expression is indicated with an orange arrow, low protein expression with a green arrow and the transitional cells are indicated with asterisks. Scale 100µm.

By P19, blastemal regions which we previously showed were positive for SIX2 and WT1 (Fig. 5F) exhibited robust expression of NCAM1, ALDH1A2, CITED1, and SIX2, shown on consecutive sections (Fig. 6C). Co-staining of consecutive sections demonstrated specific expression patterns for ALDH1A2 and CITED1, with differential expression patterns for each in distinct parts of the blastema (Fig. 6C, green arrow: high ALDH1A2, orange arrow: high CITED1, asterisk: both proteins expressed). This suggests that the blastemal-like population is a heterogenous population of cells, possibly reflecting uninduced NPC state when CITED1^high^ALDH1A2^low^, and induced NPC state when CITED1^low^ALDH1A2^high^. We found a third population which is quadruple-positive for all markers (Fig. 6C asterisks), and this could represent a transitional state between uninduced and induced blastema, which is only present in the mutant. Blastemal populations surrounded embryonic-like tubular structures, potentially remnants of UB tips. This is similar to the phenotype described by Urbach et al. (2014), who claim that these blastema structures are remains of cap mesenchymes, and that these structures are capable of differentiating into nephrogenic structures in the presence of *Wnt4* (Urbach et al., 2014). These blastemal cells remained proliferative, as evidenced by KI67 expression (Sup. Fig. 5).

## Discussion

Due to the early embryonic origin of Wilms tumors, little is known about the earliest phenotypic changes following tumor-initiating mutations. Genetically engineered mouse models provide a unique and powerful model to study these effects in a controlled and well-understood developmental context. Here we present an extensive characterization of the early events after Wilms tumor mutation induction in mouse models, which shows two patient-based Wilms tumor-initiating mutations result in fundamentally different changes to the developing kidney, and therefore different biology underlying tumors that eventually will both be classified as Wilms tumors.

We use the two previously described CSC marker sets as starting point for the detailed analysis of the direct effects of two different Wilms tumor-initiating mutations in mouse models. To enable the analysis of the Aldefluor signal using antibodies we identified ALDH1A2 as the ALDH paralogue responsible for its activity in WT CSCs (Pode-Shakked et al., 2013).(Pode-Shakked et al., 2013). Combining ALDH1A2 expression with the other WT CSC markers NCAM1, SIX2 and CITED1 we demonstrate in normal developing mouse kidneys a succession of different marker combinations in the nephrogenic lineage (Fig. 7A). An NCAM1/CITED1/SIX2 triple-positive stage in uninduced NPCs is followed by a NCAM1/ALDH1A2/SIX2-positive state in induced NPCs. As cells are epithelializing through the PTA and RV stages, SIX2 expression decreases, until only NCAM1 and ALDH1A2 remain in the comma- and S-shaped bodies. The stromal lineage is only positive for ALDH1A2 (Fig. 7A). This pattern shows that in the normal developing kidney cells are either positive for the SIX2/CITED1 CSC markers (uninduced NPCs) or the other marker set NCAM1/ALDH1A2 (comma-/S-shaped bodies) but not both. This means that either the two CSC marker sets identify different CSCs or the Wilms tumor initiating mutations force a change in CSC marker expression and induces the existence of the true Wilms tumor CSC.

**Figure 7:**
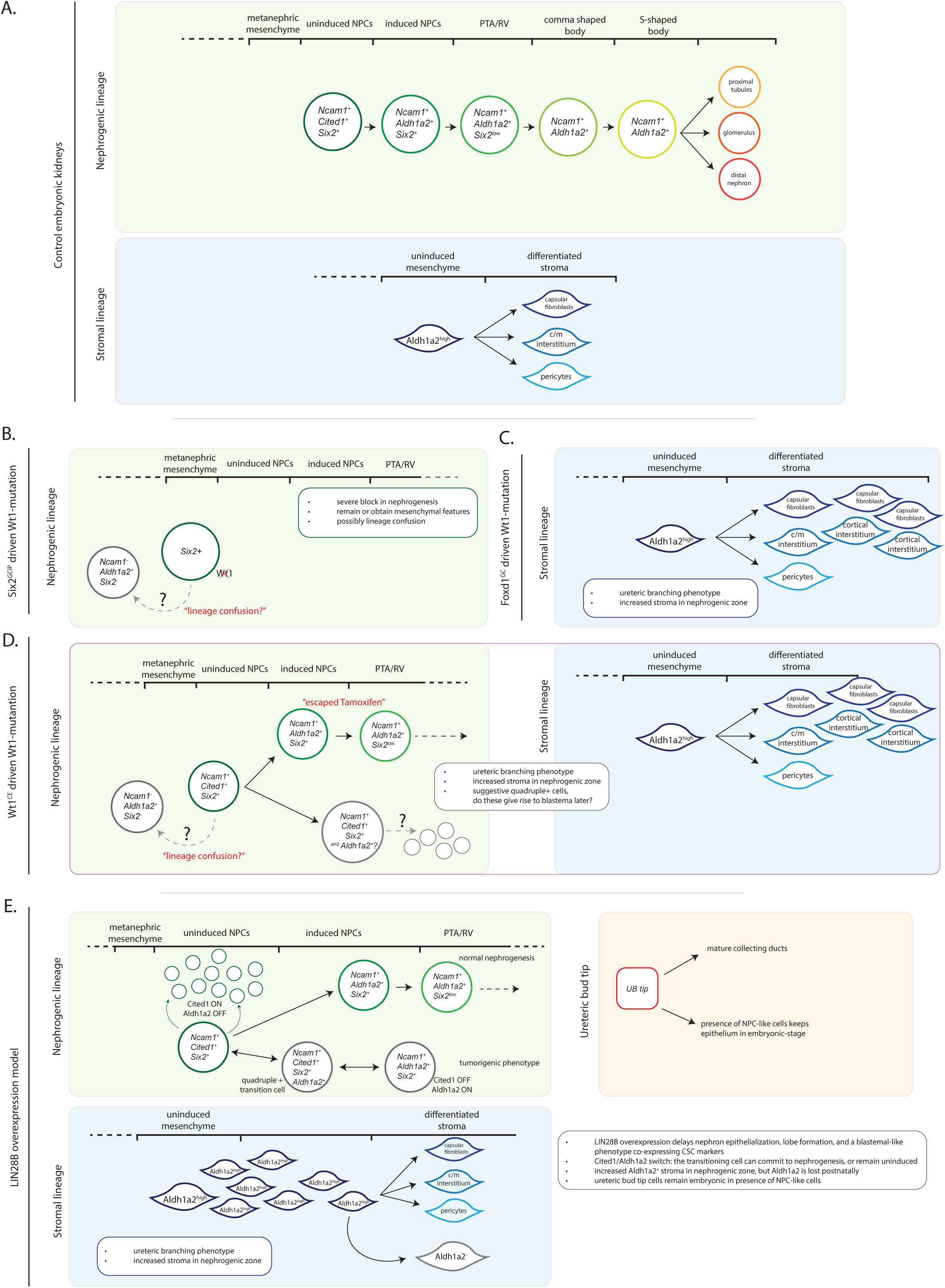
Schematic representation of genotype-phenotype speculations of Wilms tumor driver mutations. (A) CSC marker expression over the course of nephrogenesis, in the nephrogenic and stromal lineage. (B) *Six2^GCiP^* driven loss of Wt1 results in severe block of nephrogenesis, and possibly results in lineage confusion based on CSC marker expression. (C) *Foxd1^GC^* driven Wt1 loss results in expanded stroma and a ureteric bud branching phenotype. (D) Conditional loss of Wt1 through the *Wt1^CE^* tamoxifen dependent administration results in stromal and ureteric bud phenotype and a mild block of nephrogenesis. Here the first suggestive quadruple-positive CSC markers come together in the same cells possibly giving rise to blastema. (E) LIN28B overexpression results in delayed nephron epithelialization, embryonic lobe formation and blastemal-like cells co-expressing the four CSC markers.

The effect of *Wt1* loss depends on the lineage(s) where the mutation is induced. When *Wt1* was lost in the nephrogenic lineage using the *Six2*^GC^ driver we observed a very severe block in nephron formation, and instead a retention or even increase of mesenchymal appearance and behavior of the mutant cells (Fig. 7B). ALDH1A2 is the only CSC marker found in the mutant cells, which are derived from Six2^+^ cells as shown by the tdTomato-positive lineage and therefore from the nephrogenic lineage. In normal developing kidneys ALDH1A2 expression in the nephrogenic lineage is only found together with NCAM1, and in the absence of SIX2 only in the post-MET stages. The mutant kidneys do not reach this stage, so this is unlikely to explain the observed CSC marker expression. Instead, ALDH1A2^+^-only cells are a characteristic of the stromal lineage. We propose that loss of WT1 at the earliest SIX2-positive stage in the nephrogenic lineage result in a developmental shift from nephrogenic to stromal lineage, which we refer to as ‘lineage confusion’. A precedent for this is found in the *Six2*-Cre driven loss of *Pax2* which shows a comparable nephrogenic-to-stromal shift (Naiman et al., 2017).

Our analysis of stromal-specific loss of *Wt1* using the *Foxd1*^GC^ driver confirmed a previously described phenotype for this model (Fig. 7C; Weiss et al., 2020).(Fig. 7C; Weiss et al., 2020). This includes a branching phenotype in the ureteric bud and increased stroma in the nephrogenic zone. No changes in the expression pattern of the CSC markers was found. We found SIX2^+^ caps were occasionally displaced inward, raising the possibility of aberrant lobulation (see below). Important for the role of *WT1* as Wilms tumor gene, in this mutant no changes in the CSC expression patterns were found.

To knockout *Wt1* in both nephrogenic and stromal lineage we empirically maximized the efficiency of the *Wt1*^CE^ driver for combined effect in these two lineages. Using this to knockout *Wt1* we found a complex phenotype reminiscent of the previously described for the *Nes*-Cre driven *Wt1* knockout (Fig. 7D; Berry et al., 2015). This included increased stroma in the nephrogenic zone, finger-like aberrant branching of the ureteric bud and escaping nephrons that remain *Wt1*-positive, albeit fewer than in the *Nes*-Cre driven mutant. The stromal expansion in this model is much more severe than in the stromal-only *Wt1* knockout. We propose the stromal expansion found in the *Wt1*^CE^-driven mutant is a combination of a direct effect of WT1 loss in the stroma as found in the *Foxd1*^GC^-driven mutant, with the lineage confusion in the nephrogenic lineage as found in the *Six2*^GCiP^-driven model. In the E18.5 samples of the mutant we see the appearance of potential NCAM1/ALDH1A2/SIX2/CITED1 quadruple-positive cells and the first signs of clusters of blastemal cells as found in Wilms tumors. Although the host antibody incompatibility of the CITED1 and ALDH1A2 antibodies stops of from proving this quadruply positive state, it is consistent with the hypothesis stated above that the Wilms tumor initiating mutation induces the existence of the CSCs.

There are three non-mutually exclusive ways to explain why the nephrogenic phenotype in the combined nephrogenic/stromal *Wt1* knockout is less severe than in the nephrogenic-only mutant. First, we cannot determine the exact moment *Wt1* is lost in either mutant. The tdTomato gives an indication for this, but for the phenotype the (unknown) stabilities of existing Wt1 mRNA and protein can delay the moment the phenotype is induced. Second, in the *Wt1*^CE^-driven mutant the stromal component is also mutant. It is possible that the mutant stroma gives different signals to the nephrogenic lineage than normal stroma, which can modify the nephrogenic phenotype. Third, the escaping nephrons in the nephrogenic/stromal double mutant could provide molecular signals that partially rescue the mutant cells of the nephrogenic lineage. It is important to realize that in patients Wilms tumors also start developing in a further normal embryonic kidney, suggesting such a mechanism could be relevant for the origin of Wilms tumors in patients as well. The reason for the incomplete Cre activity in the *WT1*^CE^ model is not clear, but the extra level of tamoxifen control might provide additional variability due to variation in local exposure, biological processing of tamoxifen, and increased variation in the exact timing of Cre activity that could result from this could be important here. It is clear however, as was for the *Nes*-Cre driver (Berry et al., 2015) that the retention of *Wt1* expression due to incomplete Cre activity is the cause of these escaping nephrons.

Whereas the loss of *Wt1* in the nephrogenic lineage shows clear signs of lineage confusion, in which the identity of the mutant cells changes, we do not find any signs of this in the LIN28B model. Instead, this model is characterized by a delay in NPC differentiation leading to NPC-like blastemal clusters (Fig. 7E). It is again in these cells that we find potential NCAM1/ALDH1A2/SIX2/CITED1 quadruple-positive cells, though also NCAM1/CITED1/SIX2 and NCAM1/ALDH1A2/SIX2 triple positive cells. In the normal induction of NPCs there is a transition from CITED1 to ALDH1A2 in NCAM1/SIX2 double positive cells, but in normal development we have not found any sign of a quadruple-positive transition state. We propose that in this mutant model the increased LIN28B expression leads to a fluid and reversible state between the induced and non-induced NPC state and the appearance of intermediate quadruple-positive CSC marker cells that show the full complement of all four CSC markers and could form the seed of the tumor.

The signs of kidney lobulation in two of the four mutants is remarkable. The mechanism driving kidney lobulation and its physiological purpose are unknown, but lobulation is normally not a feature of murine kidneys. It is suggested that human kidney lobulation occurs through striated cells, presumably of interstitial origin, that align along the radial axis of the kidney (Lindstrom et al., 2018). This is consistent with our observation of stromal cells infiltrating the nephrogenic zone, possibly “dragging” these caps to the inner parts of the cortex. Further study on human lobulation is necessary to draw any further conclusions on this observation and its relevance for Wilms tumors.

In conclusion, by focusing on two sets of Wilms tumor cancer stem cell markers in different mouse models for Wilms tumor genes we find that a quadruple-positive state for all CSC markers is only found in the mutant kidneys, suggesting the existence of the functional CSC is induced by the tumor initiating mutation. The co-expression of CITED1 and ALDH1A2 could be the minimal set of Wilms tumor CSC markers, but antibody incompatibility prevents us from testing this. Moreover, we find clear differences in the effect of losing *Wt1* or activating LIN28B on the developing kidney, with the former leading to a change in cell identity for the mutant cells and the latter a disrupted transition in the normal nephron development cascade. A different underlying biology for these two mutant situations could provide different therapeutic opportunities for tumors caused by these mutations, or even require different therapeutic approaches. Recognizing genotype-phenotype correlations like this are essential for maximizing the chance of successful preclinical trials through selection of the right patients for the right trial. Further detailed analyses of the developmental effects of other Wilms tumor mutations is required to determine if the two phenotypes described here cover the full spectrum of Wilms tumor biology or if additional primary phenotypes exist.

## Methods

### Animals

The animal care and experimental procedures were approved by the Animal Welfare Body Leiden and are in accordance with the Dutch Experiments on Animals Act and EU Directive 2010/63/EU. For timed pregnancies, males Wt1^tm2(cre/ERT2)Wtp^/J, Wt1^tm1.2Ndha^Six2^tm1(EGFP/cre)Phoh^ and Wt1^tm1.2Ndha^B6;129S4-Foxd1^tm1(GFP/Cre)Amc^/J were crossed with female Wt1^tm1.1Ndha^ to conditionally knockout Wt1. To activate the CreERT2 in the WT2 mouse, tamoxifen (Sigma: T5648) (2mg/40g body weight) dissolved in corn oil (Sigma: C8267) was administered intraperitoneally. Pregnant females received injections at E8.5, E9.5, E10.5, E11.5, or combined at E9.5 and E10.5. To conditionally activate LIN28B, male Wt1^tm1(EGFP/cre)Wtp^/J and Wt1^tm1(EGFP/cre)Wtp^/J*B6.Cg-Igs7^tm62.1(tetO-tdTomato)Hze^/J were crossed with female B6.Cg-Gt(ROSA)26Sor^tm1(rtTA*M2)Jae^B6.Cg-Col1a1^tm2(tetO-LIN28B)Gqda^. Doxycycline (Thermo Fisher Scientific, 24390-14-5) (1g/L) was provided via the drinking water to the pregnant female mouse at E0. Pregnant females were euthanized by cervical dislocation for collection of (embryonic) kidneys. Pups at P19 were euthanized by cervical dislocation for kidney collection.

### Ex vivo kidney organ culture

Embryonic kidneys were isolated from E12.5 embryos derived from the mouse model crosses described above. Kidneys were dissected and placed on a Transwell polyester membrane insert (0.4 µm pore size, Corning, 3450) in six-well plates. The lower chamber was filled with 1ml kidney culture medium to establish an air-liquid interface. The culture medium consisted of DMEM [Gibco 41966029], MEM Non-Essential Amino Acids 1x [Gibco 11140035], Penicillin-Streptomycin 5U/ml [Gibco 15070063], βMeOH 50µM [Gibco 31350010] and 10% FCS, with or without doxycycline (2µg/mL). Cultures were maintained at 37°C in 5% CO_2_. After 1hr, when kidneys had attached to the membrane, imaging was started in an on-stage incubator chamber. Images were acquired every hour for 72-96 h. When finished, kidneys were fixed in 4% paraformaldehyde (PFA) in 1xPBS for 20’, washed 3x10’ in 1xPBS, followed by whole kidney immunofluorescent staining as described below.

### Histological and immunofluorescent staining

Freshly isolated (embryonic) kidneys or whole embryos were fixed in 4% PFA and embedded in paraffin. Smaller tissues were first embedded in 2% low-melting agarose prior to paraffin embedding. 5 µm sections were made using a microtome. Slides with sections were dewaxed with xylene and rehydrated through a series of washes with 100%, 95% and 80% ethanol. Hematoxylin (Agilent, S330930-2) staining was performed through the following protocol: slides were rinsed in deionized water, followed by 3’ in hematoxylin, rinsed in deionized water, 5’ under running tap water, 12x dipped in acid ethanol, 2x1’ in tap water, 1x2’ in deionized water. For the eosin (Sigma-aldrich, HT110232) stain, slides were stained for 30” in eosin solution, followed by 1’ in 80% ethanol, 1’ in 90% ethanol, 1’ in 100% ethanol and 2x5’ in xylene before being mounted with Pertex (Histolab, 00811-EX). For immunofluorescent staining, antigen retrieval was performed in 10mM sodium citrate (J.T.Baker, 6132043) buffer with 0.05% Tween-20 (Sigma Aldrich, 274348) (pH 6.0) in a pressure cooker for 20’. Slides were blocked in 1x PBS with 2% serum for 1hr at RT. Primary antibodies were incubated o/n at 4°C, followed by washing 3x10’ in 1xPBS the next day. Secondary antibodies were incubated o/n at 4°C, followed by co-staining for DAPI (Polysciences, 09224-10) and mounting with FluorSave reagent (Merck Millipore, 345789). Primary antibodies: Wt1 (1:50; Abcam, ab89901), Six2 (1:200; Abnova, H00010736-M01), E-cadherin/Cdh1 (1:200; BD Biosciences, 610182), Ncam1 (1:40; R&D systems (Bio Techne), AF2408), Aldh1a2 (1:100; Abcam, ab96060), Cited1 (1:200; ThermoFisher, 26999-1-AP), LTL unconjugated-biotinylated (1:400; Vector laboratories, L-1320-5), tdTomato (1:100; ThermoFisher, TA150128), Ki-67 (1:200; ThermoFisher, MA5-14520). Secondary antibodies: Goat anti-Rabbit IgG Cross-Adsorbed Secondary Antibody, Alexa Fluor 647 (1:1000; Thermofisher, A32733), Goat anti-Mouse IgG1 Cross-Adsorbed Secondary Antibody, Alexa Fluor 488 (1:1000; Thermofisher, A21121), Goat anti-Mouse IgG2a Cross-Adsorbed Secondary Antibody, Alexa Fluor 568 (1:1000; Thermofisher, A21134), Streptavidin, Alexa Fluor 546 conjugate (1:1000; Thermofisher, S11225), Donkey Anti-Goat IgG H&L (Alexa Fluor® 405) preadsorbed (1:1000; Abcam, ab175665), Donkey Anti-Rabbit IgG (H+L) Cross-adsorbed, Alexa Fluor 647 (1:1000; Thermofisher, A31573), Donkey Anti-Mouse IgG (H+L) Highly Cross-Adsorbed, Alexa Fluor 488 (1:1000; Thermofisher, A21202).

Additional methods can be found in the supplementary files.

## Supplementary

### Supplementary methods

#### Ncam1/ALDERED assay for flow cytometry

Embryonic kidneys from E12.5 embryos were directly dissociated with Trypsin/EDTA (0.05%) (Gibco, REF25300) for 3’ at 37°C. (Embryonic) kidneys from E18.5 and P19 were first minced into small pieces using a scalpel, prior to dissociation. After incubation, tissues were vigorously pipetted up and down to dissociate into single cells. DMEM + 10% FCS was used to inactivate the trypsin. Cells were centrifuged 5’ at 300g and resuspended in FACS buffer (PBS + 0.5% BSA + 0.04M EDTA). Clumps were removed by using a 35μm cell strainer. A small fraction was taken at this step to calibrate the flow cytometer. The rest of the single cells were stained for Ncam1 (1:125; NCAM-1/CD56 Antibody, R&D systems, AF2408) with a Donkey-anti-Goat Alexa Fluor 405 (1:1500; Abcam, ab175665), followed by the ALDERED assay kit (EMD Millipore, SCR150) according to the manufacturer’s protocol. LSRFortessa flow cytometer (BD Biosciences) was used to measure expression.

### Statistical analysis

Due to variable cytometer settings over time, raw data was normalized to paired littermate controls. The DEAB sample was used as the threshold for ALDERED expression. Normalized expression values were compared between controls and mutants from the same time point and Cre driver. Wt1-mutants: Six2-Cre driven E12.5 ctrl (n=9) vs mutant (n=7), Six2-Cre driven E18.5 ctrl (n=10) vs mutant (n=6), Foxd1-Cre driven E12.5 ctrl (n=13) vs mutant (n=4), Foxd1-Cre driven E18.5 ctrl (n=14) vs mutant (n=5), Wt1-CE driven E12.5 ctrl (n=9) vs mutant (n=8), Wt1-CE driven E18.5 ctrl (n=6) vs mutant (n=6)). LIN28B mutants: E12.5 ctrl (n=6) vs mutant (n=6) and E18.5 ctrl (n=6) vs mutant (n=6). Groups were compared using a non-parametric Mann-Whitney U test between groups.

## Author contributions

N.S.P. performed and analyzed all experiments. D.K., C.M.B, M.M.L., J.W.C.C., C.W.J.C and D.D.Ö supported in the performed experiments and contributed to the supplementary figures. N.S.P and P.H. designed the experiments. N.S.P. prepared figures. N.S.P., K.S.D. and P.H. wrote the manuscript.

## Acknowledgements

We thank all members of the Hohenstein lab for valuable discussions. We thank Jorrit Hos and Orhan Demirbacak for help with animals. The research was supported by KiKa (Children Cancer Free Foundation).

---

**Sup. Fig. 1:**
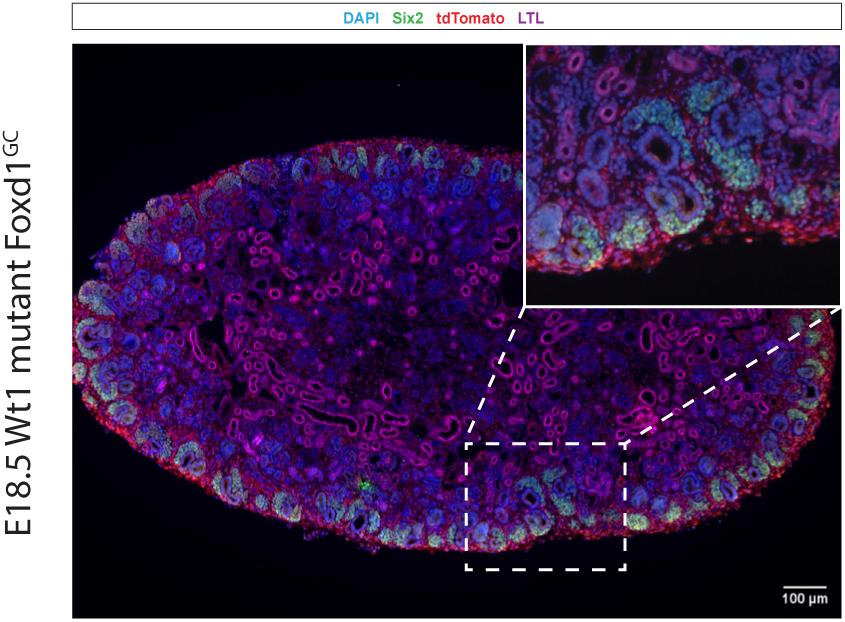
*Foxd1^GC^* driven *Wt1* mutant. SIX2+ caps lost their apical orientation and started moving inwards. Scale 100µm.

**Sup. Fig. 2:**
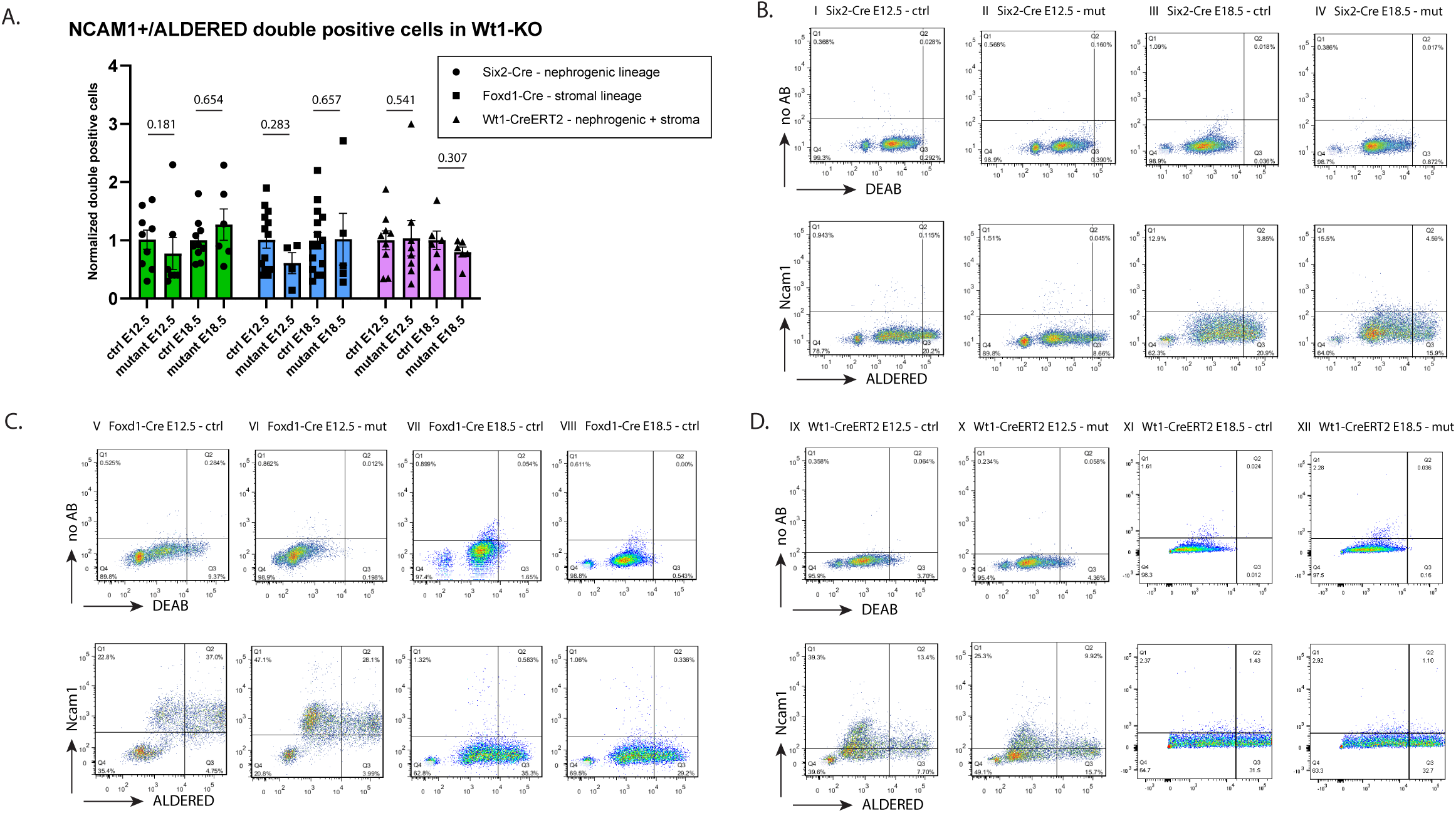
NCAM1+/ALDERED flowcytometry assay on Wt1 mutant embryonic kidneys. (A) Fraction of NCAM1+/ALDERED double positive cells in developing embryonic kidneys comparing control versus Wt1 mutants from the three lineage-specific Cre-drivers at E12.5 and E18.5. No significant differences observed between groups, using a non-parametric Mann-Whitney U test between groups. P-values indicated. (B) Raw data files from the *Six2^GCiP^* driven control versus mutant kidneys, using DEAB for setting the cutoff for the ALDH activity measured through ALDERED expression. (C) Raw data files from the *Foxd1^GC^* driven control versus mutant kidneys, using DEAB for setting the cutoff for the ALDH activity measured through ALDERED expression. (D) Raw data files from the *Wt1^CE^* driven control versus mutant kidneys, using DEAB for setting the cutoff for the ALDH activity measured through ALDERED expression.

**Sup. Fig. 3:**
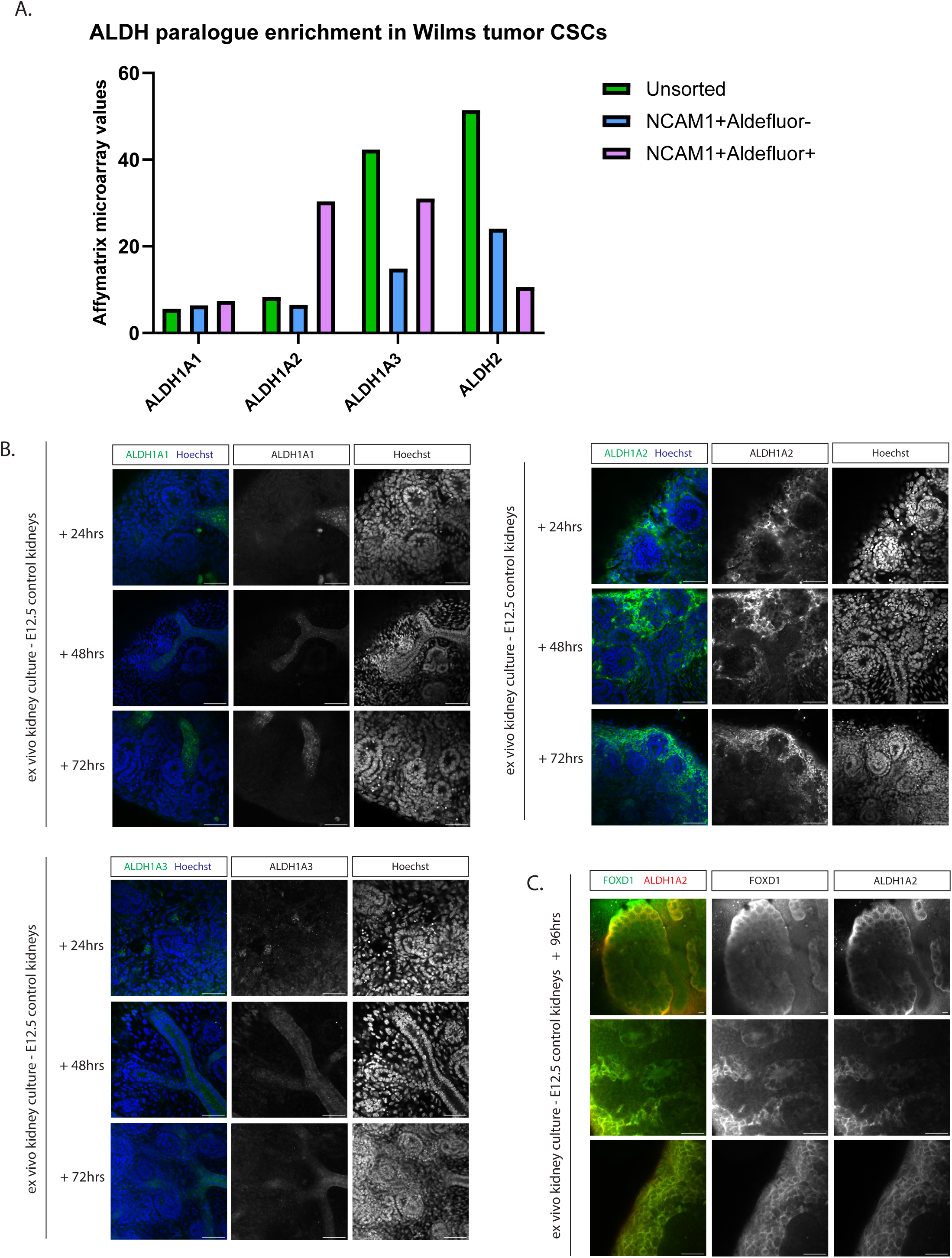
ALDH paralogue expression in Wilms tumor Xenograft microarray data and in developing kidneys. (A) ALDH paralogue enrichment in Wilms tumor Xenograft derived CSCs. A comparison between unsorted and sorted fractions, for NCAM1+/ALDEFLUOR- and NCAM1+/ALDEFLUOR+. (B) ALDH paralogue protein staining in *ex vivo* kidney cultures, cultured for 24, 48 and 72 hours. Scale bar: 50µm. (C) FOXD1 and ALDH1A2 protein expression in ex vivo kidney cultures cultured for 96 hours. Scale bar: 50µm.

**Sup. Fig. 4:**
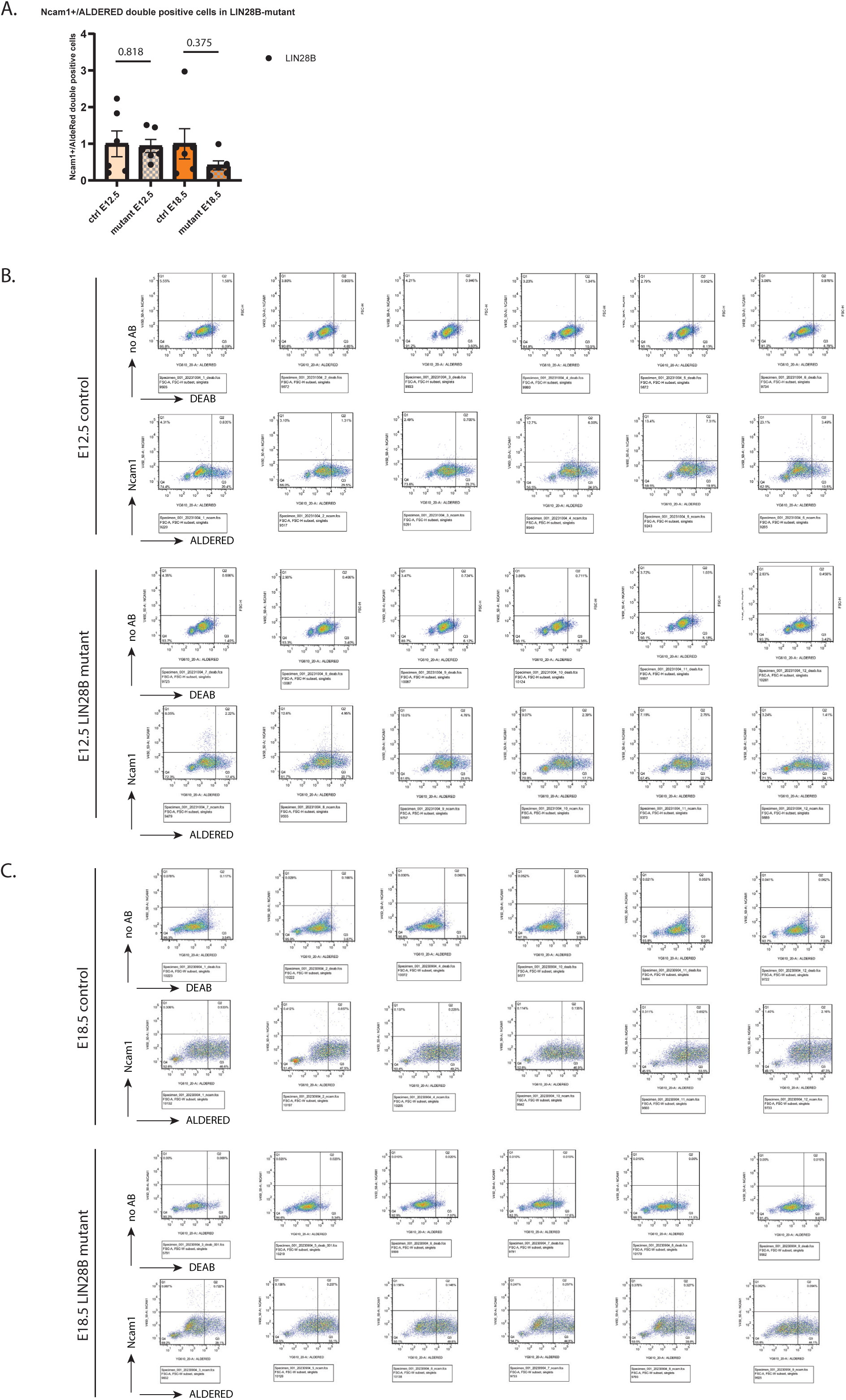
NCAM1+/ALDERED flowcytometry assay on LIN28B mutant embryonic kidneys. (A) Fraction of NCAM1+/ALDERED double positive cells in developing embryonic kidneys comparing control versus LIN28B mutants at E12.5 and E18.5. No significant differences observed between groups, using a non-parametric Mann-Whitney U test between groups. (B) Raw data files from the control versus mutant kidneys at E12.5, using DEAB for setting the cutoff for the ALDH activity measured through ALDERED expression. (C) Raw data files from the control versus mutant kidneys at E18.5, using DEAB for setting the cutoff for the ALDH activity measured through ALDERED expression.

**Sup. Fig. 5:**
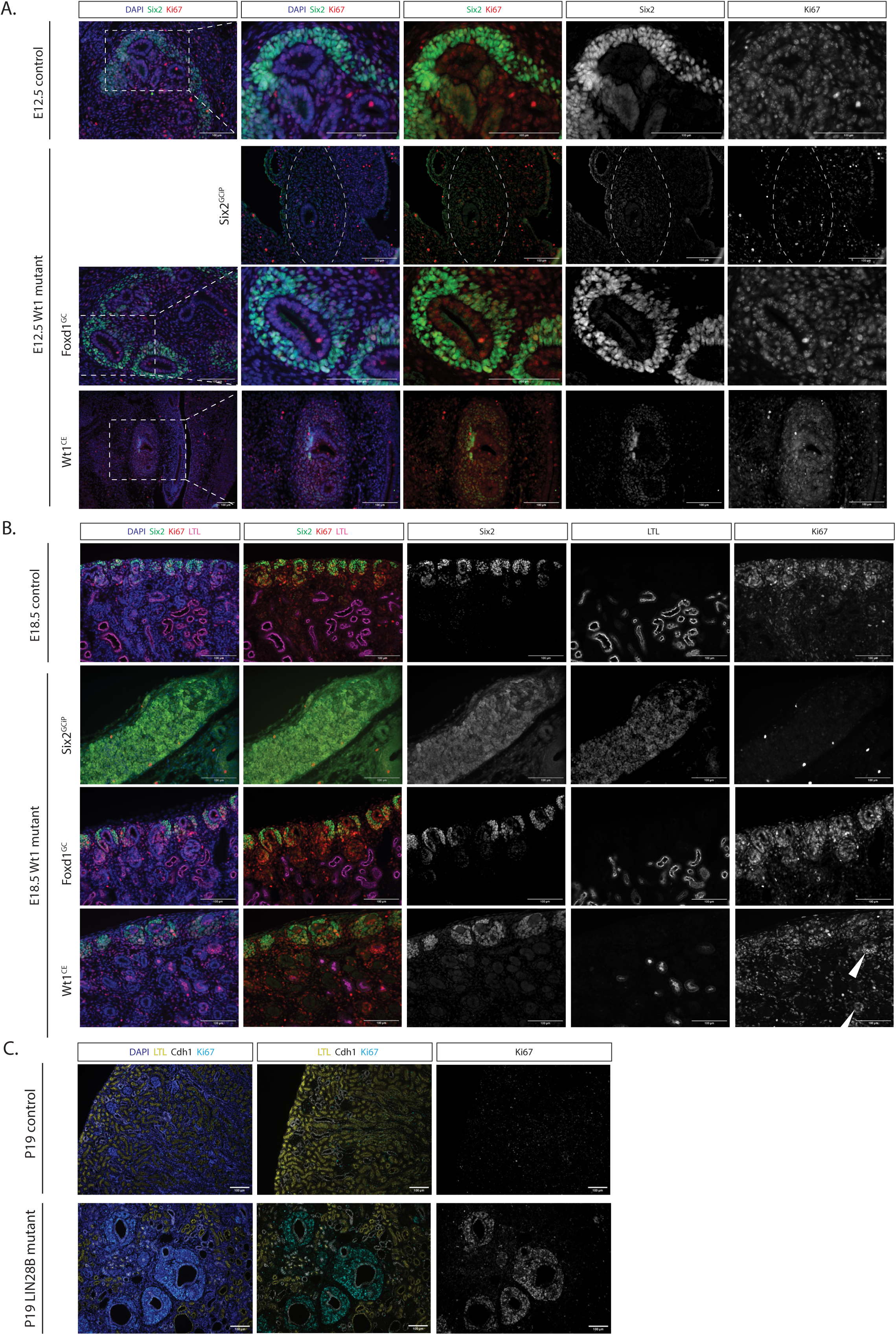
Proliferative marker expression KI67 in *Wt1* and LIN28B mutant (developing) kidneys. (A) KI67 staining for Wt1 mutant kidneys at E12.5 for lineage-specific Cre drivers. Scale 100µm. (B) KI67 staining for Wt1 mutant kidneys at E18.5 for lineage-specific Cre drivers. Scale 100µm. (C) KI67 staining for LIN28B mutant kidneys at P19. Scale 100µm.

